# MOLECULAR BASIS FOR PINK1 MATURATION

**DOI:** 10.64898/2026.08.19.745883

**Authors:** Junrong Xue, Huiqin Xu, Yanfeng Zhang, Xinrong Yu, Yan Du, Jia Guo, Jianing Duan, Weida Zhang, Xujia Liu, Yuanzhu Gao, ShuaiJiabin Chen, Sen-Fang Sui, Xiaohong Qin, Zheng Liu, Li-Zhi Mi

## Abstract

Phosphatase and tensin homolog (PTEN)-induced putative kinase 1 (PINK1), a key regulator of mitophagy, has been linked to the pathogenesis of Parkinson’s disease (PD). PINK1 recruits Parkin, an E3 ubiquitin ligase, triggering mitophagy in response to mitochondrial damage. During mitophagy, the quantity, stability, and activity of PINK1 must be strictly regulated; however, the mechanisms governing these parameters under cellular stress are still unclear. Herein, we determined the structural basis for PINK1 maturation mediated by heat shock protein 90alpha/cell division cycle 37/FK506-binding protein 51 (HSP90α/CDC37/FKBP51) chaperone complex. We identified PINK1-associated proteins using liquid chromatography– tandem mass spectrometry (LC-MS/MS) and determined the structures of the complexes using Cryo-Electron Microscopy (Cryo-EM). Results showed that FKBP51 potentially interacts with a conserved leucine–proline–phenylalanine (LPF) motif on the activation loop of PINK1 and negatively regulates PINK1 functions in mitophagy. A PINK1 mutation located at the FKBP51 recognition site is linked to mitophagy deficiency, which can be partially rescued by specific inhibition of FKBP51. These findings reveal a general mechanism for PINK1 recognition by the HSP90α/CDC37/FKBP51 chaperone complex and suggest a potential approach for upregulating PINK1 activity, which is impaired in PD.

## INTRODUCTION

Mitophagy is a strictly regulated process that removes damaged mitochondria with a depolarized membrane potential or overloaded calcium^1,2^. Mitophagy dysregulation disrupts mitochondrial bioenergetics, leading to the release of reactive oxygen species and pro-apoptotic factors from damaged mitochondria^1,2^. Consequently, mitophagy dysregulation is often associated with neurodegenerative diseases, inflammation, and cardiovascular and metabolic diseases.

PINK1, a mitochondrial kinase, plays a critical role in sensing mitochondrial damage and triggering mitophagy^3–6^. Loss-of-function mutations of PINK1 are linked to the early onset of autosomal recessive Parkinson’s disease (EOPD) ^7,8^. Therefore, developing therapeutic strategies targeting PINK1-deficient PD is an essential but challenging task.

Under resting conditions, PINK1 is transported into mitochondria, where it is cleaved and degraded, maintaining a low level and activity^9^. Upon mitochondrial damage, the full-length PINK1 accumulates on the outer membrane of mitochondria (OMM) and, subsequently, gets activated through dimerization and association with the translocase of the outer membrane (TOM) complex^10–12^, leading to the recruitment of Parkin, an E3 ubiquitin (Ub) ligase, to damaged mitochondria in order to initiate mitophagy^13–15^. Precisely regulating the quantity, stability, and activity of PINK1 in sensing mitochondrial damage is crucial for coordinating this event cascade in mitophagy. However, the underlying mechanisms regulating them are still unclear.

Molecular chaperones are essential for precisely regulating the quantity and activity of many proteins related to physiology and pathology^16,17^. These chaperones, including heat shock protein 90 (HSP90), function with their cochaperones to facilitate the folding, stabilization, degradation, translocation, and interactions of client kinases in cell-specific and context-dependent manners^16–18^. Notably, PINK1 is a client kinase of HSP90^19,20^.

Among the cochaperones of HSP90, cell division cycle 37 (CDC37) is critical for the folding and stabilization of kinase clients^18,21–23^. CDC37 facilitates the transfer of client kinases from heat shock protein 70 (HSP70) to HSP90, inhibits the adenosine triphosphatase (ATPase) activity of HSP90, and stabilizes unfolded kinases within the chaperone complex^21–24^.

FK506-binding protein 51 (FKBP51) is another cochaperone for HSP90^25–27^. It is a tetratricopeptide repeat (TPR)-containing peptidyl-prolyl *cis*/*trans* isomerase (PPIase) involved in inflammation, innate immunity, stress response, and neuronal protection^25–27^. Although its functions in the activation and maturation of steroid receptors have been researched, its role in the folding, activation, and maturation of client kinases is still unclear^25,28^.

In this study, we identified PINK1 as a novel kinase client of FKBP51 and determined the molecular interactions and underlying mechanisms that enable HSP90α, CDC37, and FKBP51 to dynamically regulate the quantity, stability, and activity of PINK1 during mitophagy. Our findings will provide a framework for understanding the chaperone-mediated regulation of the activity of PINK1 during mitophagy and offer insights into developing targeted compounds for enhancing PINK1-mediated mitophagy.

## RESULTS

### PINK1 is a novel kinase client for FKBP51

To investigate how PINK1 interacts with other proteins in regulating mitophagy, we transiently transfected the full-length PINK1 into the human embryonic kidney cell line HEK293T and purified the endogenous PINK1 complexes using affinity and gel-filtration chromatography (Fig. 1a,b). The PINK1 complexes (in the second fraction) were eluted from the gel filtration column in a single, symmetric peak. We performed liquid chromatography–tandem mass spectrometry (LC-MS/MS) analysis of this peak fraction to identify PINK1-associated proteins and determined the Cryo-EM structures of these endogenous PINK1 complexes at a resolution of 2.7–2.8 Å.

**Fig. 1.**
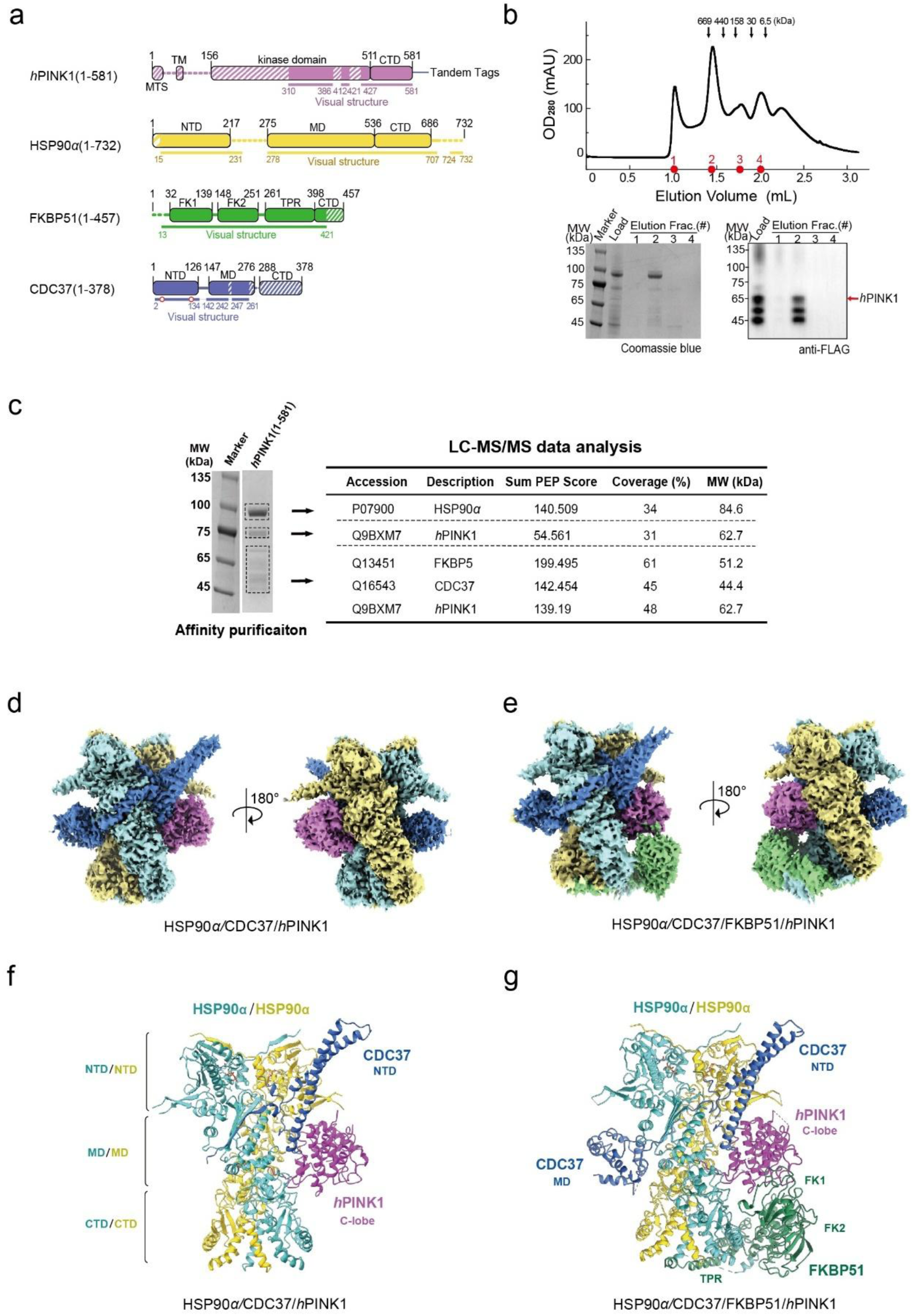
Identification of PINK1 as a client of the HSP90α/CDC37/FKBP51 complex by Cryo-EM. **(a)** Schematic of the domain organization of proteins in the PINK1-chaperone complex. Striped/dashed regions denote areas lacking interpretable density. Red circles indicate phosphorylated residues (pS13 and pT118). **(b)** Size-exclusion chromatography profile of the affinity-purified PINK1 complex. Elution fractions were analyzed by SDS-PAGE with Coomassie blue staining (lower left) and Western blotting (lower right). **(c)** Mass spectrometry identification of the protein bands. **(d/e)** Cryo-EM densities of the PINK1/HSP90α/CDC37 complex (d) and the PINK1/HSP90α/CDC37/FKBP51 complex (e). **(f/g)** Structural overviews of the PINK1/HSP90α/CDC37 complex (f) and the PINK1/HSP90α/CDC37/FKBP51 complex (g).

LC-MS/MS analysis identified HSP90α, CDC37, and FKBP51 as PINK1-associated proteins (Fig. 1c). Their Cryo-EM structures revealed two distinct HSP90α/CDC37– PINK1 assemblies, one containing a bound FKBP51 and the other without (Fig. 1d– g, Supplementary Figs. 1 and 2, and Table 1).

**Fig. 2.**
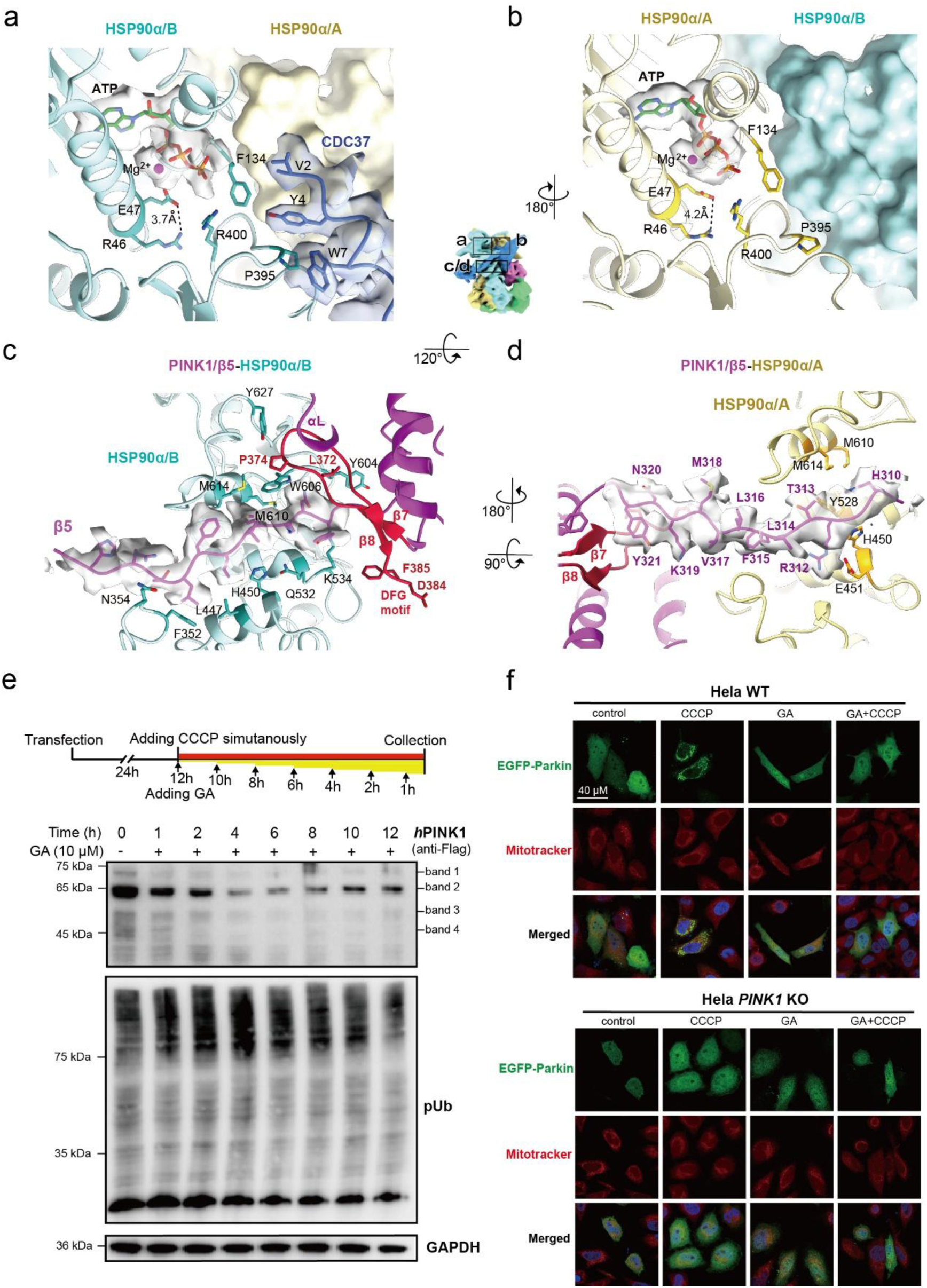
Stabilization of inactive PINK1 by HSP90α. (a/b) ATP-binding sites in HSP90α subunit A (a) and subunit B (b). ATP densities are shown as transparent surfaces. Subunit A is colored yellow and subunit B cyan. **(c/d)** The PINK1 binding interface on HSP90α subunit B (c) and subunit A (d). Density for the PINK1 β5 strand is contoured at 3σ and displayed as a transparent gray surface. **(e)** Time-course analysis of PINK1 stability and activity in transfected HEK293T cells treated with 5 μM CCCP and/or 10 μM geldanamycin (GA). Top, treatment scheme; bottom, representative Western blot. **(f)** Representative confocal images showing PINK1-mediated recruitment of EGFP-Parkin to mitochondria in WT and *PINK1*-KO HeLa cells under treatment with 5 μM CCCP, 2 μM geldanamycin, both, or none.

In both assemblies, two HSP90α proteins functioned as a central organizer, coordinating the interactions of the protomers within the complex. The two HSP90α proteins adapted in a closed conformation, forming an elongated, pseudo-symmetric dimer (Fig. 1f,g). PINK1 was partially unfolded, with its N-lobe disordered and its β5 strand embedded within the lumen of the HSP90α dimer. The N-terminal domain of CDC37 interacted with PINK1 and one HSP90α protomer, while its M domain bound to the other HSP90α protomer (Fig. 1g).

In the FKBP51-bound assembly (FKBP51/HSP90α/CDC37–PINK1), the FKBP51 TPR domain bound to the HSP90α C-terminal dimeric interface (Fig. 1g). The FK1 domain of FKBP51 interacted with the C-lobe of PINK1. However, the FK2 domain did not directly interact with other proteins in the complex (Fig. 1g).

To validate our structural models, we immunoprecipitated endogenous PINK1 from WT HeLa cells and subjected the associated complexes to Blue Native PAGE analysis. Endogenous PINK1 formed a large complex with HSP90α, FKBP51, and TOM20 upon CCCP treatment (Supplementary Fig. 3a). This suggests that the chaperone-bound PINK1 pool may dynamically exchange with the mitochondrial import machinery, or that a fraction of PINK1 escapes chaperone-mediated retention^29^. Co-immunoprecipitation further confirmed the endogenous PINK1-FKBP51 interaction in both HEK293T and WT HeLa cells (Supplementary Fig. 3b). In summary, these results suggest that PINK1 can function as a novel kinase client for FKBP51.

### HSP90α serves as a scaffold to stabilize partially folded PINK1 and prevent its degradation

PINK1 recognition by CDC37 and HSP90α was mediated by highly conserved interfaces (Supplementary Fig. 3c and Supplementary Video S1), suggesting these interactions are crucial for the physiological functions of PINK1.

As a carrier of kinase clients, CDC37 delivered partially folded PINK1 to HSP90α. The N-terminal residues of CDC37, including V2, Y4, and W7, fit into a hydrophobic cavity of HSP90α (Fig. 2a). In addition, phosphorylated S13 stabilized the orientation of the HPNI fragment of CDC37 (amino acids 13–34; Supplementary Fig. 3d–f). As such, the HPNI fragment could be inserted into a highly conserved pocket in the PINK1 C-lobe close to the β7 and β8 strands (Supplementary Fig. 3e,f). This HPNI fragment mimicked the configuration of the correctly folded PINK1 αi–β4 loop, thereby stabilizing the PINK1 C-lobe structure (Supplementary Fig. 3e,f). The density of CDC37’s M domain was not well resolved as it dynamically interacted with the disordered PINK1 N-lobe, as visualized in our three-dimensional variability analysis (3DVA; Supplementary Video S2).

Two HSP90α proteins adapted in a closed conformation, with their N-terminal domains engaged in an adenosine triphosphate (ATP)-bound state (Fig. 2a,b). Arginine 46 (R46) formed a salt bridge contact with the catalytic residue E47 (Fig. 2a,b), promoting E47 deprotonation. This salt bridge contact is proposed as a critical switch for coupling ATP hydrolysis to the global conformational changes in HSP90^30^.

Like other kinase clients of HSP90, the β5 strand of PINK1, which is essential for the folding and assembly of functional PINK1, was stretched and embedded in a tightly packed hydrophobic tunnel formed by the M domains of the two HSP90α proteins (Fig. 2c,d, Supplementary Fig. 4a–c, and Supplementary Video S3). Leucine 316 (L316) of PINK1 was anchored at the center of this tunnel. This central hydrophobic residue was highly conserved across the PINK1 family, while the flanking polar residues were variable (Supplementary Fig. 4c,d). Mutations of the hydrophobic residues along the β5 strand diminished the HSP90α–PINK1 association during immunoprecipitation (Supplementary Fig. 4e). Moreover, in the reported active PINK1 structures, the β5 strand was required for organizing the hydrogen-bonding network of the N-lobe of PINK1 (Supplementary Fig. 3e and Supplementary Video S3) ^12,31^. If the β5 strand was retracted from this network, the entire hydrogen-bonding network necessary for the proper folding of PINK1 was destroyed. As a consequence, the density of the N-lobe of PINK1 was not resolvable in our structures.

HSP90α–PINK1 binding also changed the conformation of the C-terminal domain (CTD) of PINK1, blocking the kinase assembly. Specifically, HSP90α–PINK1 binding disrupted the secondary structure of the αL helix and displaced the αK helix of the PINK1 CTD (Supplementary Fig. 3e). Since the αK helix is essential for the association of the N-terminal αA helix of PINK1 with the CTD during PINK1 importation, its displacement impaired the assembly and activation of PINK1 (Supplementary Fig. 3e) ^32^. Notably, five of seven PD-associated mutations located at the PINK1 CTD were enriched on this αK helix (Supplementary Fig. 4c) ^32^.

Moreover, compared with RAF1- and CDK4-bound HSP90αs, HSP90α–PINK1 binding induced conformational changes in HSP90α such that the α15’ of HSP90α became disordered, creating a hydrophobic cavity to accommodate the β7–β8 loop of PINK1 (Supplementary Fig. 4f). Three PD-associated mutations have been identified in the β7 strand of PINK1, with one of them (I368N) being reported to lower PINK1 levels and impair the mitochondrial importation of PINK1 under stress^33^.

Using geldanamycin to specifically inhibit the ATPase activity of HSP90α in transfected HEK293T cells promoted PINK1 degradation in a time-dependent manner. Compared to the sample treated with only 5 μM CCCP, adding 10 μM geldanamycin for 12 hours led to >80% enhancement in PINK1 degradation under CCCP-induced mitochondrial depolarization (Fig. 2e and Supplementary Fig. 4g). However, geldanamycin alone only had a mild effect on the Ub phosphorylation activity of PINK1 (Fig. 2e and Supplementary Fig. 4h).

Inhibition of HSP90α also downregulated PINK1-mediated Parkin recruitment under CCCP-induced mitochondrial depolarization. Compared to the sample treated with only 5 μM CCCP, adding 2 μM geldanamycin diminished the recruitment of EGFP-Parkin to mitochondria in WT HeLa cells (Fig. 2f). This CCCP-induced mitochondrial recruitment of EGFP-Parkin was abolished in *PINK1-* knockout HeLa cells, confirming its PINK1 dependence (Fig. 2f).

### FKBP51 dynamically interacts with the PINK1 activation loop to negatively regulate PINK1 activity

In the HSP90α/CDC37/FKBP51–PINK1 assembly, FKBP51 was anchored to the C-terminal dimeric interface of HSP90α via its TPR domain (Fig. 3a,b and Supplementary Video S4). The α7E helix from the FKBP51 TPR domain bound to the hydrophobic groove formed at the dimeric interface of the HSP90α CTDs (Supplementary Fig. 5a). The helical symmetry of the α7E helix was broken at alanine 410 (Ala410) to accommodate its mismatch with the C2 symmetry of the HSP90α dimer (Supplementary Fig. 5a). The C-terminal tail of the HSP90α subunit B formed an L-shaped clamp to secure the α7E helix of FKBP51 in place (Supplementary Fig. 5a). In addition, the conserved methionine–glutamate– glutamate–valine–aspartate (MEEVD) motif from HSP90α subunit B formed a short helix embedded into the cavity of the FKBP51 TPR domain (Supplementary Fig. 5b).

**Fig. 3.**
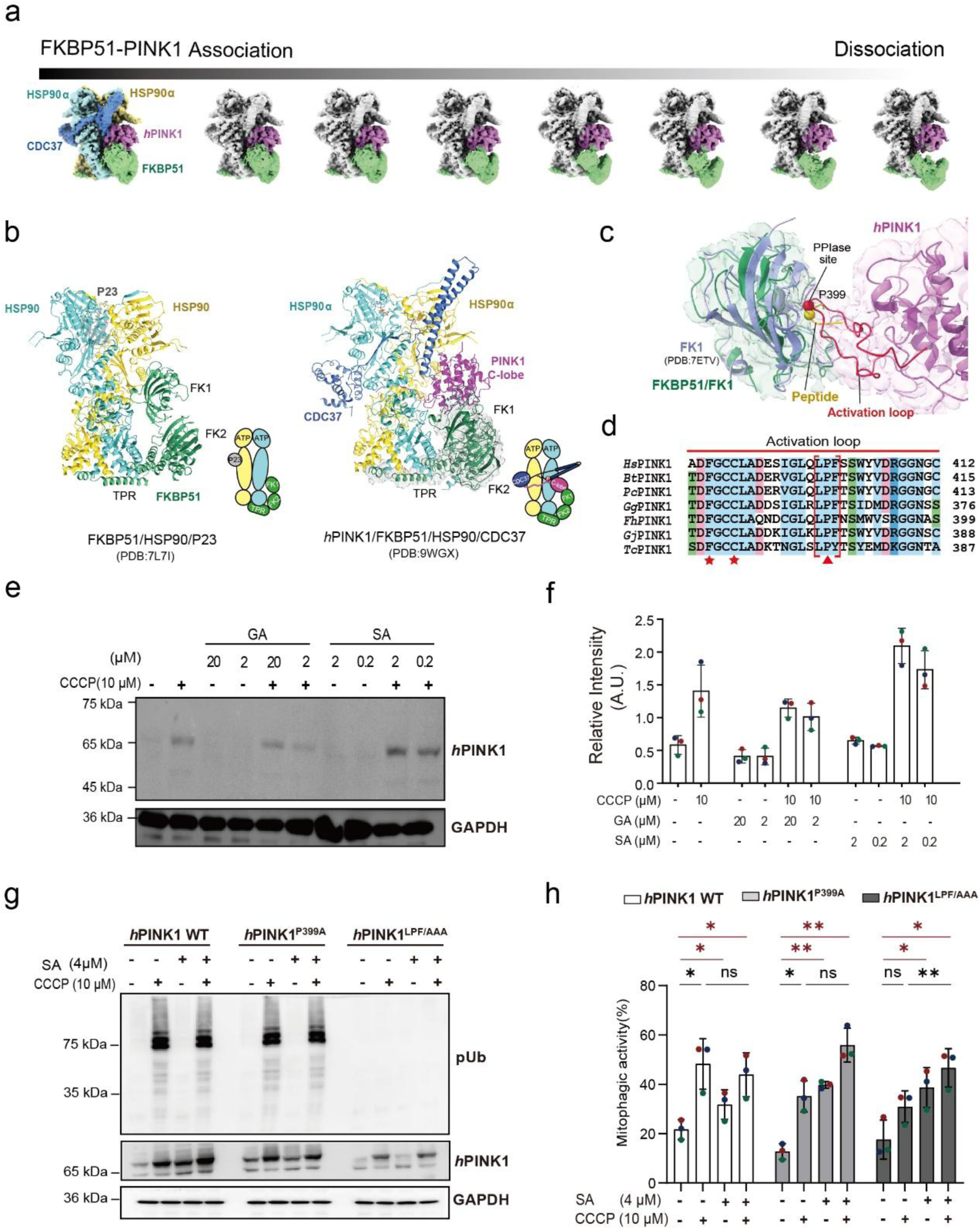
Regulation and recognition of PINK1 by FKBP51. **(a)** 3D variability analysis of dynamic FKBP51-CDC37 interactions with PINK1 in the chaperone complex. Classified density maps are segmented and displayed in surface representation. **(b)** Structural comparison of the HSP90-FKBP51 complex with and without PINK1. **(c)** Interaction of the FK1 domain of FKBP51 with the PINK1 activation loop. The FKBP51-PINK1 complex is shown in cartoon representation with the overlaid transparent local density map (low-pass filtered to 5 Å, contoured at 3σ). The FKBP51-peptide structure (PDB 7ETV) is superimposed for comparison. The PINK1 activation loop is colored red and the FK1-bound peptide is colored yellow. Prolines recognized by FK1 are shown as spheres. **(d)** Sequence alignment of the PINK1 activation loop. PD-associated mutations are marked with red stars or triangles; the conserved LPF/Y motif is highlighted by red brackets; the proline at the FKBP51 isomerase recognition site is indicated by a red triangle. **(e/f)** Expression and stabilization of endogenous PINK1 in HeLa cells treated overnight with the indicated concentrations of CCCP, GA, or SA. Representative Western blot (e) and quantification from triplicate experiments (f). **(g)** Effect of FKBP51 Inhibition on Ubiquitin phosphorylation by WT PINK1 and mutants. *PINK1*-KO COS7 cells transfected with PINK1^WT^, PINK1^P399A^ or PINK1^LPF/AAA^ were treated with SAFit2 and/or CCCP. Ub phosphorylation was analyzed by Western blotting in triplicate. **(h)** Mitophagy activity of PINK1 ^WT^, PINK1^P399A^ and PINK1^LPF/AAA^ upon SAFit treatment under mitochondrial depolarization. *PINK1*-KO HeLa cells were co-transfected with plasmids encoding PINK1 ^WT^, PINK1^P399A^ and PINK1^LPF/AAA^ together with mt-Keima. Cells were treated with or without SAFit2 and/or CCCP for 10 h. Mitophagy was analyzed by FACS as previously described ^50^. The statistics are determined from triplicate biological repeats.

Upon FKBP51*–*PINK1 binding, the anchored FKBP51 underwent conformational changes (Fig. 3b). A comparison of the FKBP51 conformations in the FKBP51/HSP90/P23 (PDB 7L7I)^34^ and FKBP51/HSP90α/CDC37–PINK1 complexes showed that the individual TPR, FK1, and FK2 domains were nearly identical. However, the linkers connecting these individual domains varied between these two structures^34^. This resulted in a 19° rotation of the FK1 domain and a 7° rotation of the FK2 domain relative to the FK2 and TPR domains, respectively (Fig. 3b).

Along with these conformational changes, FKBP51 dynamically interacted with PINK1, in conjunction with CDC37 (Fig. 3a). Our 3DVA results showed that FKBP51 lifting was coupled with the twisting of the N-terminal double helices of CDC37 during their interactions with PINK1 (Supplementary Video S4). In this dynamic process, the α7E helix of FKBP51 served as a pivot for lifting FKBP51, while the MEEVD motif of HSP90α acted as a latch that secured the positioning of the FKBP51 TPR domain (Supplementary Fig. 5a,b).

FKBP51 used its PPIase-competent FK1 domain to interact with PINK1. Due to the dynamic nature of FKBP51–PINK1 interactions, the density of their binding interface was not well resolved in the entire complex structure. To study the FKBP51–PINK1 interactions in detail, we calculated a local density map for FKBP51 touching PINK1 at 3.59 Å resolution (Supplementary Fig. 5c–f). Using this map lowpass-filtered at 5 Å resolution, we could trace the entire backbone of the PINK1 activation loop loop as polyalanine, except for glycine residues (Fig. 3c). Further refinement of this model against the 3.59 Å map (Table 1) showed that the traced activation loop was in proximity to the FKBP51 active site, indicating that FKBP51 might use its FK1 domain to catalyze the isomerization of a specific proline on the PINK1 activation loop or stabilize the activation loop in a specific conformation. Indeed, there is a conserved LPF motif on the PINK1 activation loop, while the proline 399 (P399) mutation on this motif has been identified in a patient with PD (Fig. 3d) ^35^. When the PINK1- and peptide-bound FKBP51 structures (PDB 7ETV)^36^ were superimposed, the Cα–Cα distance between two prolines was 1.5 Å, indicating that the conserved LPF motif on the PINK1 activation loop was positioned roughly at the PPIase active site of FKBP51 (Fig. 3c,d and Supplementary Fig. 5g). Consistently, fluorescent anisotropy assay revealed markedly reduced binding of the LPF/AAA peptide to the purified FK1 domain of FKBP51 relative to peptides containing the LPF or LAF motifs across all concentrations tested (Supplementary Fig. 5h). Moreover, the conformation of the PINK1 activation loop in FKBP51-bound PINK1 resembled that of the inactive Src kinase but not that of the active PINK1 predicted using Google DeepMind AlphaFold^31^ (Supplementary Fig. 5i).

To validate our structural model, we used SAFit2, a specific PPIase inhibitor of FKBP51, to study the functions of FKBP51 in regulating PINK1 activity. Unlike geldanamycin, 10 μM SAFit2 had a minimal effect on PINK1 degradation in transfected HEK293T cells treated with 5 μM CCCP (Supplementary Fig. 6a–c). In addition, adding SAFit2 did not affect the Ub phosphorylation activity of PINK1 under CCCP-induced mitochondrial depolarization. However, adding SAFit2 increased the expression and/or stabilization of the full-length endogenous PINK1 in WT HeLa cells treated with 10 μM CCCP overnight (Fig. 3e,f).

Importantly, mutation of the FK1-recognizable LPF motif on the PINK1 activation loop impaired PINK1-mediated activities during mitophagy^36^. We introduced two mutations (PINK1^P399A^ and PINK1^LPF/AAA^) on the PINK1 activation loop and analyzed their effects on PINK1-mediated Ub phosphorylation and mitophagy under CCCP-induced mitochondrial depolarization. Results showed that the PINK1^LPF/AAA^ mutation, but not the PINK1^P399A^ mutation, significantly reduced specific activities of PINK1 under CCCP-induced mitochondrial depolarization (Fig. 3g and Supplementary Fig. 6d).

Moreover, the PINK1^P399L^ mutation, which has been identified in patients with PD with unknown pathological consequences, also impaired the Ub phosphorylation activity of PINK1 (Supplementary Fig. 6e) ^35^. However, SAFit2 had no detectable impact on the Ub phosphorylation activity of WT PINK1 and its mutants in the presence or absence of CCCP (Fig. 3g and Supplementary Fig. 6d,e). In mitophagy studies, the PINK1^LPF/AAA^ mutation impaired PINK1-mediated mitophagy under CCCP-induced mitochondrial depolarization. Inhibition of FKBP51 by SAFit2 promoted mitophagy in cells expressing WT PINK1 or its mutants, even without CCCP-induced mitochondrial depolarization. As such, adding SAFit2 rescued the impaired mitophagy activity of the PINK1^LPF/AAA^ mutant under CCCP-induced mitochondrial depolarization (Fig. 3h and Supplementary Fig. 6f,g).

In summary, these results suggest that FKBP51 can potentially interact with the LPF motif on the PINK1 activation loop. FKBP51 is involved in the regulation of PINK1-mediated mitophagy, either by stabilizing a specific conformation of the PINK1 activation loop or by catalyzing the *cis*/*trans* isomerization of P399. FKBP51 inhibition can rescue the impaired mitophagy activity of a specific PINK1 mutant, PINK1^LPF/AAA^, at the PINK1 activation loop.

## DISCUSSION

### Principal findings

Our study clarifies a link between chaperone-mediated PINK1 folding/unfolding and PINK1 activity regulation under mitochondrial stress (Fig. 4). Under normal physiological conditions, the newly synthesized PINK1 is unfolded for transportation into and out of mitochondria via the TOM complex^11^. Misfolded PINK1 is degraded, ensuring that PINK1 quantity and activity are kept low to protect healthy mitochondria^37,38^. Upon mitochondrial damage, PINK1 transportation via the TOM complex is blocked, leading to the accumulation and activation of full-length PINK1 on the OMM. Activated PINK1 recruits Parkin and subsequently promotes clearance of damaged mitochondria using autophagosomes^10,11^. The HSP90/CDC37 complex is a key regulator of PINK1 stability, degradation, and importation^19,20,33,39^. However, the exact mechanisms governing HSP90/CDC37–PINK1 interactions, as well as the assembly and functions of the PINK1–chaperone complex, are still unclear. In this study, we found that PINK1 forms a large complex with endogenous HSP90α, CDC37, and FKBP51. However, HSP90α and FKBP51 have different functions in regulating PINK1 functions. HSP90α and CDC37 bind to partially unfolded PINK1 for either PINK1 stabilization or mitochondrial transportation, so geldanamycin-induced specific inhibition of HSP90α leads to the degradation of both full-length as well as presenilin-associated rhomboid-like protein/mitochondrial processing peptidase (PARL/MPP)-cleaved PINK1 in a time-dependent manner. In contrast, FKBP51 has a marginal effect on PINK1 stabilization; rather, FKBP51 contributes to downregulating the mitophagy activity of PINK1 in the absence of CCCP-induced mitochondrial depolarization (Fig. 3h). These functions of the HSP90α/CDC37/FKBP51 chaperone complex establish a link between PINK1 folding and its stability, activity, and transportation in mitophagy. This link enables cells to respond to cellular and environmental stress, thereby retaining mitochondrial integrity and promoting the clearance of damaged mitochondria for overall cellular health.

**Fig. 4.**
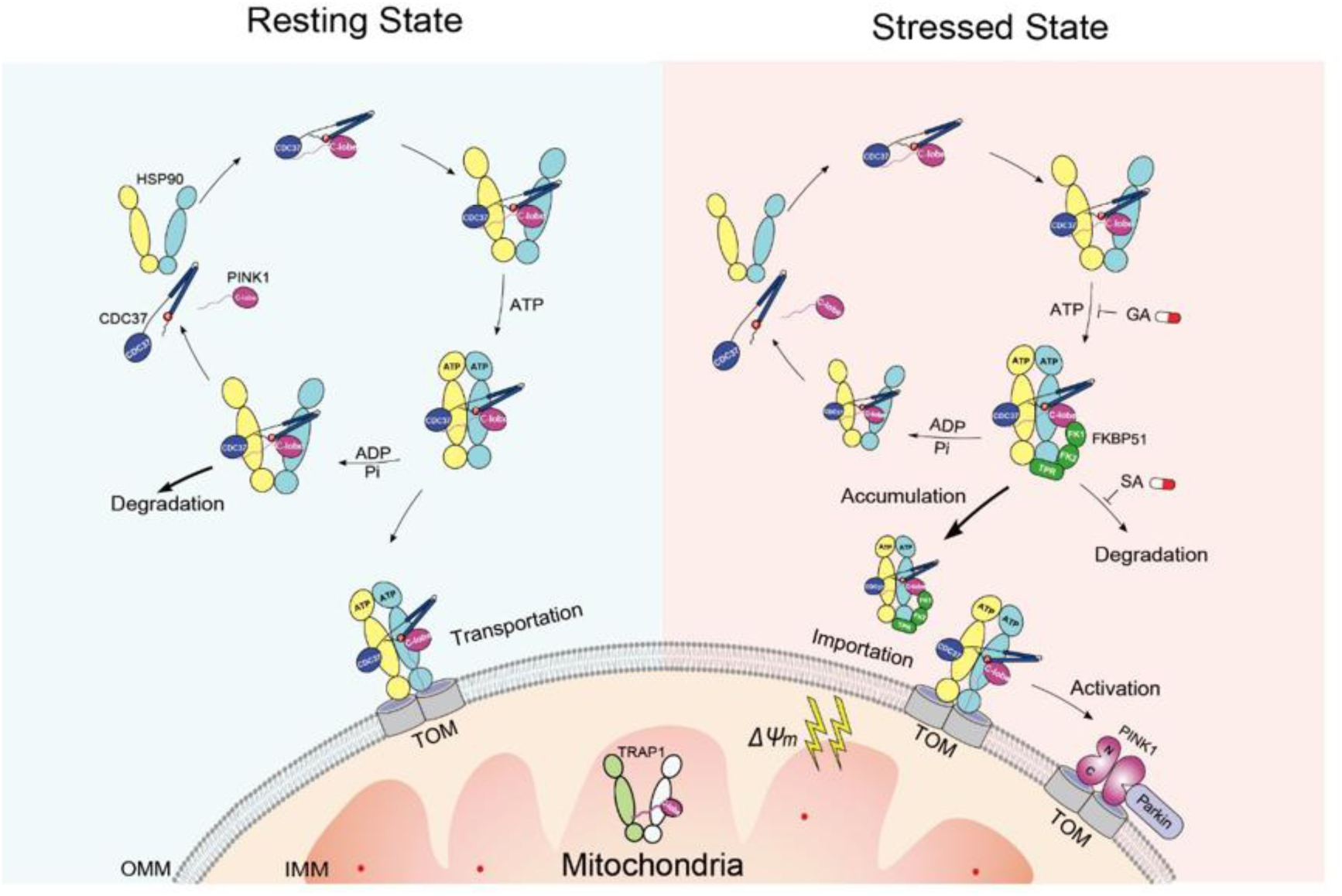
Model for PINK1 regulation by the HSP90α/CDC37/FKBP51 chaperone machinery under resting and stressed conditions. Under normal physiological conditions, newly synthesized unfolded PINK1 is stabilized by the HSP90/CDC37 complex and subsequently imported into the mitochondria. Following cleavage by PARL at the inner mitochondrial membrane, PINK1 is retro-translocated to cytosol and degraded by the ubiquitin–proteasome system. This regulatory mechanism prevents uncontrolled PINK1-mediated mitophagy (left). Under stress conditions, upregulation of HSP90α and FKBP51 promotes the formation of a complex with CDC37-bound PINK1. As import of PINK1 into damaged mitochondria is blocked, PINK1 accumulates and is activated through dimerization, thereby recruiting Parkin to initiate mitophagy. Notably, the HSP90 inhibitor geldanamycin suppresses PINK1 folding and maturation, leading to its degradation. Conversely, FKBP51 negatively regulates PINK1 activity during mitochondrial clearance. Specific inhibition of FKBP51 by SAFit2 enhances PINK1 maturation, promoting the clearance of damaged mitochondria (right).

### Implications

Our study provides a framework for understanding the molecular mechanisms underlying the recognition and regulation of kinase clients by the HSP90/CDC37/FKBP51 chaperone complex. To date, it was unclear how FKBP51 recognizes and/or alters the conformations of kinase clients. In previously reported PINK1 structures, the activation loop is well ordered and extended, adopting an active conformation suitable for substrate binding^40,41^. In contrast, in our Cryo-EM structures, the conformation of the PINK1 activation loop resembled that of inactive Src kinase and was positioned near the FKBP51 active site (Fig. 3c and Supplementary Fig. 5i) ^42^. As the PINK1 activation loop contains a conserved LPF motif, we investigated whether FKBP51 recognized this motif and how this recognition is involved in regulating PINK1 activity. P399L mutation within this LPF motif is linked to PD pathology and results in impaired PINK1 activity in Ub phosphorylation^35^. Similarly, LPF/AAA mutation diminishes PINK1 activity in Ub phosphorylation and mitophagy under CCCP-induced mitochondrial depolarization. Inhibition of FKBP51 by SAFit2 can rescue the impaired mitophagy activity of the PINK1^LPF/AAA^ mutant under mitochondrial depolarization. These functional data validate that FKBP51 can recognize the PINK1 activation loop for negative regulation of PINK1 activity in mitophagy. In this process, FKBP51 can presumably catalyze *trans*-to-*cis* isomerization of P399 on the PINK1 activation loop or stabilize the loop in its inactive conformation. Aligned with the LPF motif of PINK1, cyclin-dependent kinase 4 (CDK4) contains a conserved TPV motif in its activation loop (Supplementary Fig. 4c). FKBP51 catalyzes proline isomerization in this TPV motif to inhibit CDK4 activation, suggesting such recognition mode might be applicable to other kinase clients of FKBP51^43^. However, the low resolution (5**–**6 Å) and structural heterogeneity surrounding the PINK1 activation loop precluded precise delineation of the molecular basis by which FKBP51 recognizes and regulates PINK1 conformational transitions. Further studies are therefore warranted to elucidate the underlying mechanism.

In our structures, FKBP51 was anchored to the HSP90α dimeric interface through its TPR domain to interact with PINK1. Our MS analysis also identified several other TPR domain–containing proteins, such as TOMM70 and FKBP52, which could potentially compete with FKBP51 in binding to HSP90α and thus interplay with FKBP51 in regulating PINK1 folding, degradation, translocation, or interactions in mitophagy. In addition, SAFit2 may affect mitophagy by inhibiting FKBP51 functions independently of mitochondrial depolarization^44^. Therefore, the detailed mechanisms governing the FKBP51-induced downregulation of PINK1 mitophagy activity are inevitably complicated and should be clarified in future studies. Nevertheless, our findings offer valuable insights into the recognition of kinase clients by FKBP51 and the pathogenesis of PD-associated mutations.

It is now increasingly important to develop targeted therapeutics for diseases related to impaired mitophagy. In developing PINK1-specific therapeutics, pioneering teams focus on enhancing PINK1 activity using ATP analogs^45^. However, such strategies are still under debate^46,47^. Our study found that FKBP51 can negatively regulate PINK1-mediated mitophagy in a manner independent of CCCP-induced mitochondrial depolarization (Fig. 3e–j). Accordingly, specific inhibition of FKBP51 can rescue the impaired mitophagy activity of the PINK1^LPF/AAA^ mutant, demonstrated as proof-of-concept in our experiments. Consistent with our findings, FKBP51 has been actively explored as a druggable target for treating stress-related diseases^26,48^. Moreover, SAFit2 is a potential compound for treating neuroinflammation, depression, neuropathic pain, and Huntington’s disease^26,49^. Therefore, our study highlights an alternative approach for intervention in PD and mitochondria-related diseases.

### Conclusions

The Cryo-EM structures of the HSP90α/CDC37/FKBP51–PINK1 complex revealed molecular interactions that enable HSP90α, CDC37, and FKBP51 to dynamically regulate the quantity, stability, activity, and transportation of PINK1 during mitophagy. A mutation on the PINK1 activation loop, which is associated with PD, is located proximally to the FKBP51 active site. Mutations at this site obstruct PINK1 functions in mitophagy, which can be partially rescued by specific inhibition of FKBP51. In summary, our findings provide a foundation for understanding the structure of PINK1 and its recognition and regulation by the HSP90/CDC37/FKBP51 chaperone complex. They also provide insights into alternative intervention strategies for PD and mitochondria-related diseases.

## METHODS

### Plasmid construction

The gene encoding *h*PINK1 (residues 1–581) was subcloned into the pEF-1 vector at the *BamH*I and *Xho*I sites to generate a C-terminal 3×FLAG–Strep II fusion. Plasmids encoding *h*PINK1^C377A^ (residues 1–581) and *h*PINK1^C549A^ (residues 1–581) were generated similarly. The EGFP-Parkin expressing plasmid was constructed as previously described.

For co-expression studies, WT *h*PINK1 (residues 1–581) and EGFP-Parkin were cloned into the pCDH vector by homologous recombination. This vector contains a dual-promoter cassette: *h*PINK1 is driven by the CMV promoter and EGFP-Parkin by the EF-1α promoter. Plasmids for co-expression of *h*PINK1^P399A^ or *h*PINK1^LPF/AAA^ (residues 1–581) with EGFP-Parkin were constructed analogously.

### Expression and purification of *h*PINK1 (1-581)

HEK293T cells were cultured in 15-cm dishes to ∼80% confluency. For transfection, 75 μg plasmid DNA and 225 μg PEI were each dissolved in PBS. The PEI solution was added dropwise to the plasmid solution, mixed gently, and incubated at room temperature for 10 minutes before addition to the cell culture media. After 5 h, the medium was replaced with fresh medium. Cells were harvested 48 h post-transfection.

The culture medium was removed, and cells were resuspended in 2 mL PBS. Cell suspensions were centrifuged at 1,000 × *g* for 5-10 min at 4°C, and the supernatant was discarded. Cell pellets were flash-frozen in liquid nitrogen and stored at -80°C.

Frozen cells were thawed and resuspended in lysis buffer (20 mM HEPES, pH 7.5; 300 mM NaCl; 10% glycerol; 1 mM PMSF; 0.5% Triton X-100; 1 mM EDTA). The suspension was passed through a 20-gauge needle 20-30 times and centrifuged at 20,000 × *g* for 30 min at 4°C to obtain cleared lysate.

Cleared supernatant was incubated with Strep II-affinity resin with gentle shaking for 1 h. The resin was loaded onto a column and washed with 30 column volumes (CV) of Wash Buffer I (20 mM HEPES, pH 7.5; 150 mM NaCl; 10% glycerol; 1 mM PMSF; 0.02% Triton X-100), followed by 30 CV of Wash Buffer II (20 mM HEPES, pH 7.5; 150 mM NaCl; 0.02% Triton X-100). Bound proteins were eluted using Elution Buffer (20 mM HEPES, pH 7.5; 100 mM NaCl; 0.01% Digitonin; 5 mM biotin).

Affinity-purified samples were further purified by size-exclusion chromatography on a Superose 6 increase 3.2/300 column (Cytiva) equilibrated with 20 mM HEPES (pH 7.5), 100 mM NaCl, 0.01% Digitonin, and 1 mM DTT.

### Analysis of mass spectrometry data

Protein bands excised from Coomassie blue-stained SDS-PAGE gels were digested with 0.5 µM trypsin for 16 h. The resulting peptides were analyzed by nano-flow liquid chromatography-tandem mass spectrometry (nLC-MS/MS) using a an EASY-nLC 1200 system coupled to an Orbitrap Fusion Lumos mass spectrometer (Thermo Fisher Scientific).

Peptides were loaded onto a 150 μm × 2 cm self-packed C18 trap column (3 μm particle size; Dr. Maisch GmbH, Germany) and separated on a 150 μm × 30 cm self-packed C18 analytical column (1.9 μm particle size; Dr. Maisch GmbH). Mobile phase A consisted of 0.1% formic acid in water; mobile phase B consisted of 0.1% formic acid in 80% acetonitrile. The gradient for mobile phase B was: 8-12% over 10 min, 12-27% over 69 min, 27-45% over 28 min, 45-95% over 3 min, followed by a 10-min hold at 95%.

The mass spectrometer was operated in data-dependent acquisition (DDA) mode using Xcalibur 4.0 software. Full-scan MS spectra were acquired in the Orbitrap (m/z 350–1800) at a resolution of 120,000. MS2 spectra were generated by higher-energy collisional dissociation (HCD) at a normalized collision energy of 30% and acquired in the ion trap.

MS/MS spectra were searched against the UniProt Swiss-Prot human database (release August 2018, 20,325 entries) using Proteome Discoverer (Version 2.2; Thermo Fisher Scientific). Search criteria included: full tryptic specificity, two missed cleavages allowed, carbamidomethylation as a fixed modification, and oxidation as a dynamic modification. Precursor ion mass tolerance was set to 20 ppm for all MS in the Orbitrap, and fragment ion mass tolerance was 0.6 Da for all MS2 spectra in the ion trap. A high confidence score filter (FDR < 1%) was applied to select target peptides, and their corresponding MS/MS spectra were manually inspected.

### Cryo-EM sample preparation and data collection

For cryo-EM sample preparation, 4 μL of the *h*PINK1(1–581)-chaperone complex was applied to glow-discharged 300-mesh UltrAuFoil (2/1) grids. Grids were blotted for 4 s at 4 °C and 100% humidity before plunge-freezing into liquid ethane using a Vitrobot Mark IV (Thermo Fisher Scientific).

Grid screening and data collection were performed on a 300-kV Titan Krios microscope (Thermo Fisher Scientific) equipped with a K3 Summit direct electron detector and a GIF Quantum energy filter. Movie stacks were automatically collected using EPU in super-resolution mode at a calibrated magnification corresponding to 0.668 Å per physical pixel, with a target defocus range of -1.2 to -2.0 μm. The energy filter slit width was set to 20 eV, and the total electron dose was 60 e^-^/Å² for each micrograph stack.

### Cryo-EM image processing

Images were processed with cryoSPARC^51^ and RELION^52^. The processing strategies were shown in Supplementary Fig. 1. Movie stacks were motion-corrected using MotionCor2^53^, and contrast transfer function (CTF) parameters were estimated by Patch CTF. Micrographs with visible contaminations or an maximum resolution worse than 5 Å were excluded, yielding 19,051 micrographs for further processing.

Particles was auto-picked from 200 micrographs using BLOB picker and classified into 50 classes via 2D classification. Representative 2D class averages were used to train Topaz^54^ for automated particle picking. Reference-free 2D classification yielded a stack of 1,989,484 particles, which were subjected to *ab initio* reconstruction followed by heterogeneous refinement. Particles yielding high-resolution maps with well-defined features were selected for homogeneous refinement.

Selected particles underwent 3D classification in RELION, and representative classes were refined by Non-Uniform refinement in cryoSPARC. This yielded 3D reconstructions at 2.79 Å for the *h*PINK1/HSP90α/CDC37/FKBP51 complex and 2.67 Å for the *h*PINK1/HSP90α/CDC37 complex^52,55^. After particle subtraction and local refinement with a mask encompassing FKBP51 and PINK1, a local map of the FKBP51-PINK1 complex was obtained at 3.59 Å^52,55^.

All reported resolutions were estimated according to the gold-standard FSC 0.143 criterion. Local-resolution distributions were calculated using ResMap^56^. Three-dimentional FSC maps and model Q-scores were calculated as described previously^57,58^.

### Model building and refinement

Initial models were generated by docking the HSP90 complex (PDB ID 5FWM) and AlphaFold2-predicted *h*PINK1 into cryo-EM density maps using UCSF Chimera^31,59^. Models were iteratively rebuilt in COOT and refined in Phenix^60,61^. The geometry of the refined models was validated using MolProbity^62^. Data collection and refinement statistics are reported in Supplemental Table 1. Coordinates and cryo-EM maps have been deposited in the PDB and EMDB, respectively.

### Inhibition of HSP90 ATPase activity by geldanamycin

HEK293T cells were cultured in DMEM supplemented with 10% FBS at 37°C in a CO₂ incubator. At ∼80% confluency, cells in 10-cm dishes were transfected with 25 μg plasmid encoding FLAG-tagged full-length *h*PINK1 using 75 μg PEI in PBS. After 5 h, the medium was replaced and cells were reseeded into a 24-well plate.

After 36 h, 5 μM CCCP was added, followed by 10 μM geldanamycin to triplicate wells at the indicated time points (1–12 h). Cells were harvested after 12 h of CCCP exposure, washed with 500 μL PBS and lysed in 80 μL RIPA buffer supplemented with 1 mM PMSF and 1 mM Na₃VO₄. Lysates were frozen at -20°C for ≥30 min, thawed and cleared by centrifugation at 12,000 rpm for 10 min at 4°C. Supernatants were mixed with 6× SDS loading buffer for Western blot analysis.

Proteins were separated by 8% SDS–PAGE and transferred to PVDF membranes. Membranes were blocked with 5% non-fat dry milk in TBST for 1 h and incubated overnight at 4°C with anti-FLAG antibody. After washing with TBST, membranes were incubated with horseradish peroxidase-conjugated goat anti-rabbit IgG (Abmart Cat #: M21002) for 1 h at 4°C.

Western blot images were acquired using an LAS-4000 imager (GE Health), and band intensities were quantified using the manufacturer’s software. Statistics were calculated from triplicate technical replicates.

### Inhibition of FKBP51 PPIase activity by SAFit2

Inhibit of FKBP51 PPIase activity were performed as described above for HSP90 ATPase inhibition, except that SAFit2 (10 μM) was substituted for geldanamycin (10 μM).

### Parkin recruitment to mitochondria

HeLa cells (wild-type or *h*PINK1-KO) were cultured in DMEM supplemented with 10% FBS in 35-mm confocal dishes at 37°C in a CO₂ incubator. Before transfection, the medium was replaced with fresh medium. At ∼80% confluence, cells were transiently transfected with 2.5 μg EGFP-Parkin plasmid using 7.5 μg PEI per well. After 4 h, the medium was replaced.

After 24 h, cells were treated for 1 h with 5 μM CCCP, 2 μM geldanamycin, both, or left untreated, in the presence of 100 nM MitoTracker Deep Red FM (Yeasen Biotechnology). Cells were then fixed and stained with 10 μg/mL DAPI. Images were acquired using a Leica SP8 confocal microscope equipped with an HC PL APO CS2 63×/1.40 oil-immersion objective. DAPI, EGFP-Parkin, and MitoTracker signals were detected sequentially using 405-nm, 488-nm, and 638-nm laser lines, respectively.

### Analysis of PD-associated mutations on PINK1-mediated Parkin recruitment

*h*PINK1-KO HeLa cells were cultured in DMEM supplemented with 10% FBS in 35-mm confocal dishes at 37°C in a CO₂ incubator. Just before transfection, the culture medium was replaced with fresh medium. At ∼80% confluence, cells were transiently transfected with bicistronic plasmids encoding *h*PINK1 WT, *h*PINK1^P399A^ or *h*PINK1^LPF/AAA^ together with EGFP-Parkin using PEI. After 4 h, the medium was replaced.

After 24 h, cells were treated for 1 h with 5 µM CCCP, 4 μM SAFit2, both, or left untreated, in the presence of 100 nM MitoTracker Deep Red FM (Yeasen Biotechnology). Cells were fixed and stained with 10 μg/mL DAPI. Images were acquired using a Leica SP8 confocal microscope equipped with an HC PL APO CS2 63×/1.40 oil-immersion objective at 23°C. DAPI, EGFP-Parkin, and MitoTracker signals were detected sequentially using 405-nm, 488-nm and 638-nm laser lines, respectively.

### Endogenous PINK1 stability assays

WT HeLa cells were seeded in 24-well plates and cultured in DMEM supplemented with 10% FBS at 37°C in a CO₂ incubator. At ∼80% confluence, cells were treated for 12 h with the indicated combinations of CCCP (10 μM), geldanamycin (2 or 20 μM), and/or SAFit2 (200 nM or 2 μM), or left untreated.

After treatment, cells were washed with 500 μL PBS and lysed in 80 μL RIPA buffer supplemented with 1 mM PMSF. Lysates were frozen at -20°C for ≥30 min, then thawed and cleared by centrifugation at 12,000 rpm for 10 min at 4°C. Supernatants were mixed with 6× SDS loading buffer for Western blot analysis.

Proteins were separated by 8% SDS-PAGE and transferred to PVDF membranes. Membranes were blocked with 5% non-fat dry milk in TBST for 1 h and incubated overnight at 4°C with anti-PINK1 antibody (Cell Signaling Technology, Cat #: 6946, 1:500 dilution). After washing with TBST, membranes were incubated with horseradish peroxidase-conjugated goat anti-rabbit IgG (Abmart, Cat #: M21002) for 1 h at 4°C.

Western blot images were acquired using an LAS-4000 imager (GE Healthcare), and band intensities were quantified using ImageQuant LAS 4000 software (GE Healthcare). Statistics were calculated from triplicate technical replicates.

### Expression and purification of the FK1 domain of FKBP51

The gene encoding the FK1 domain of FKBP51 with a C-terminal MBP tag was cloned into the pET-28a vector. The plasmid was transformed into *E. coli* Rosetta (DE3) competent cells. A single colony was inoculated into 1 L LB medium and cultured at 37℃ until the OD_600_ reached 0.6. Protein expression was induced with 0.6 mM IPTG at 37℃ for 4 h. Cells were harvested by centrifugation and lysed by sonication in 40 mL lysis buffer (20 mM Tris-HCl, pH 8.0). The lysate was cleared by ultracentrifugation at 18,000 rpm for 50 min. Cleared lysate was loaded onto a Ni-NTA column (YEASEN, 20502ES60). The column was washed with wash buffer (50 mM HEPES, pH 8.0, 200 mM NaCl, 30 mM Imidazole) and bound proteins were eluted with 10 mL elution buffer (50 mM HEPES, pH 8.0, 200 mM NaCl, 300 mM Imidazole). Eluted proteins were collected and concentrated using a 3-kDa-cutoff centrifugal concentrator and further purified by gel-filtration chromatography on a Superdex 75 Increase 10/300 GL column (Cytiva, 29148721). Purity was assessed by Coomassie blue-stained SDS-PAGE.

### Fluorescent anisotropy binding assays

FITC-PEG_2_-LQLAFSS, FITC-PEG_2_-LQAAASS and FITC-PEG_2_-LQLPFSS (>95% purity) were synthesized by Chutai Biotechnology (Shanghai, China), dissolved in PBS, aliquoted as 1 mM stock solutions and stored at -80 ℃. purified FK1 domain was serially diluted twofold. FITC-labeled peptide was added to FK1 at the indicated concentrations to a final concentration of 10 nM in a total volume of 200 µL. After mixing, samples were transferred into a 96-well plate (Beyotime, FCP966) and fluorescent anisotropy was measured on an EnSpire Multilabel Reader (PerkinElmer) with PBS as a blank. Data are from three biological replicates, each with three technical replicates.

### Blue Native PAGE

The Blue native PAGE was performed as previously described^11^. The following buffers were used: cathode buffer B (50 mM Tricine, 7.5 mM Imidazole, 0.02% Coomassie blue G-250, pH 7.0), cathode buffer B/10 (50 mM Tricine, 7.5 mM Imidazole, 0.002% Coomassie blue G-250, pH 7.0), and anode buffer (25 mM Imidazole pH 7.0). Electrophoresis was initiated at 100 V with a current limit of 300 mA. After 18 min, the voltage was increased to 500 V and the current limit was reduced to 15 mA. When the running front reached one-third of the gel length, cathode buffer B was replaced with cathode buffer B/10. Electrophoresis was continued until the front reached the bottom of the gel.

Proteins were transferred to PVDF membrane and detected by Western blotting using antibodies against FKBP51(Proteintech, cat #67874-1-Ig), TOM20 (MedChemExpress, cat # YA1380), *h*PINK1 (Cell Signaling Technology, cat # 6946S), and HSP90α (Proteintech, cat # 13171-1-AP).

### Flow cytometry analysis of mitophagy

*PINK1*-KO Hela cells were cultured with DMEM supplemented with 10% FBS at 37 ℃ in a CO_2_ incubator. Plasmids encoding *h*PINK1^WT^, *h*PINK1^P399A^ or *h*PINK1^LPF/AAA^were co-transfected with mt-Keima into *PINK1*-KO Hela cells using Lipo8000 (Beyotime). After 24h, the medium was replaced, and cells were treated for 10 h with 10 µM CCCP, 4 µM SAFit2, both, or left untreated. Cells were then subjected to flow cytometry as previously described ^63,64^. The statistics were determined from triplicated biological replicates.

### Western blot analysis of Ubiquitin phosphorylation by *h*PINK1

*PINK1*-KO COS7 cells were seeded in 12-well plates and cultured in DMEM supplemented with 10% FBS at 37 °C in a CO_2_ incubator. Cells were transfected with plasmids encoding *h*PINK1^WT^, *h*PINK1^P399A^ and *h*PINK1^LPF/AAA^. After 24 h, the medium was replaced, and cells were treated for 3 h with 10 µM CCCP, 4 µM SAFit2, both, or left untreated.

Cells were washed twice with PBS and lysed in 100 µL/well lysis buffer (20 mM Tris-HCl, pH 8.0, 150 mM NaCl, 1 mM EDTA, 1 mM EGTA, 1% NP-40, 1% Sodium deoxycholate, 2.5 mM Sodium pyrophosphate, 1 mM beta-glycerophosphate, 1 mM Na_3_VO_4_, 1µg/mL leupeptin, 1 mM PMSF, and 1x complete protease inhibitor cocktail (Roche)). Lysate were cleared by centrifugation and mixed with 6× SDS loading buffer. Proteins were separated by reducing SDS-PAGE and transferred to PVDF membranes. Membranes were probed with antibodies against PINK1, phosphorylated Ubiquitin (pUb) and GAPDH. Band intensities were quantified using ImageJ. Statistics was calculated from triplicated biological replicates.

### Immunoprecipitation of endogenous *h*PINK1-associated proteins

WT HeLa cells were cultured in DMEM supplemented with 10% FBS at 37 ℃ in a CO_2_ incubator. Cells were seeded in 10-cm dishes and, at ∼80% confluence, treated with or without 10 µM CCCP for 8 h. Cells were then scraped, collected and washed three times with PBS. Cell pellets were resuspended in 700 µL lysis buffer (20 mM HEPES, pH 7.5, 300 mM NaCl, 10% Glycerol, 1 mM PMSF, 0.5% Triton X-100, 1 mM EDTA). Lysates were passed through a 20-gauge needle 10 times and cleared by centrifugation at 16,000 *g* for 5 min at 4 ℃.

Cleared supernatant was incubated with 5 µL anti-PINK1 antibody and 50 µL protein A beads at 4 ℃ for 5 h. Beads were washed twice with 400 µL lysis buffer. Bound proteins were eluted with 100 µL elution buffer (50 mM Glycine, pH 2.7, 1 mM PMSF, 1 mM Na_3_VO_4_) and immediately neutralized with 40 µL 1 M Tris-HCl (pH 8.0). Samples were analyzed by Blue Native PAGE and Western blotting using antibodies against *h*PINK1, HSP90α, FKBP51, and TOM20.

### Co-immunoprecipitation of endogenous *h*PINK1-associated FKBP51

WT HeLa cells were cultured in DMEM supplemented with 10% FBS at 37 ℃ in a CO_2_ incubator. Cells were seeded in six 10-cm dishes. At ∼80% confluence, cells in three dishes were treated with 10 µM CCCP for 8 h, while the remaining three were left untreated. Cells were then scraped, collected and washed three times with PBS. Cell pellets were resuspended in 700 µL lysis buffer (20 mM HEPES, pH 7.5, 300 mM NaCl, 10% Glycerol, 1 mM PMSF, 0.5% Triton X-100, 1 mM EDTA). Lysates were passed through a 20-gauge needle 10 times and cleared by centrifugation at 16,000 *g* for 5 min at 4 ℃.

Cleared supernatant was incubated with 5 µL anti-PINK1 antibody and 50 µL protein A beads at 4 ℃ for 5 h. Beads were washed twice with 400 µL lysis buffer. Bound proteins were eluted with 100 µL elution buffer (50 mM Glycine, pH 2.7, 1 mM PMSF, 1 mM Na_3_VO_4_) and immediately neutralized with 40 µL 1 M Tris-HCl (pH 8.0). Samples were analyzed by SDS PAGE and Western blotting using an antibody against FKBP51 (Santa Cruz. Cat #, sc-271547).

Co-immunoprecipitation of endogenous *h*PINK1-associated FKBP51 from HEK293T cells was performed analogously.

### Immunoprecipitation analysis of *h*PINK1-HSP90α association

*PINK1*-KO Hela cells were cultured in DMEM supplemented with 10% FBS at 37 ℃ in a CO_2_ incubator. At ∼80% confluence, cells were transfected with plasmids encoding *h*PINK1^WT^, *h*PINK1^F315E^, *h*PINK1^L316E^*, h*PINK1^M318E^ or *h*PINK1^P374E^ using Lipo8000 (Beyotime Biotechnology). After 24 h, the medium was replaced. After a further 12 h, cells were treated with 10 µM CCCP for 8 h. Cells were scraped, collected and centrifuged at 16,000 *g* for 5 min. Cell pellets were resuspended in 300 µL lysis buffer (50 mM Tris-HCl, pH 7.4, 150 mM NaCl, 10% Glycerol, 1 mM PMSF, 0.5% Triton X-100, 1 mM EDTA, 1mM Na_3_VO_4_, 1× Protease inhibitor cocktail from Roche). Lysate were passed through a 20-gauge needle 10 times and cleared by centrifugation at 16,000 *g* for 5 min.

Cleared supernatant was incubated with 3 µL anti-HSP90α antibody and 30 µL protein A beads at 4 ℃ for 5 h. Beads were washed twice with 200 µL lysis buffer. Bound proteins were eluted with 50 µL elution buffer (50 mM Glycine, pH 2.7) and immediately neutralized with 20 µL 1 M Tris-HCl (pH 8.0). Samples were analyzed by SDS-PAGE and Western blotting using antibodies against *h*PINK1, HSP90α, and GAPDH.

### Quantifications and statistical analysis

For Western blotting, images were acquired using an LAS-4000 imager (GE Health), and band intensities were quantified using the manufacturer’s software. Means and standard deviations were calculated from triplicate experiments. Statistical significance was assessed using a paired T-test on the results from technical replicates (ns = not significant; *, p < 0.05; **, p < 0.01; ***, p < 0.001; ****, p < 0.0001). Data were processed and plotted using OriginPro (OriginLab).

For flow cytometry analysis, gating strategies are shown in Supplementary Fig. 6h,I. Mitophagy activity was quantified as the percentage of mt-Keima-positive cells with a 561 nm/405 nm fluorescence intensity ratio above the gated threshold. Data are from triplicated biological replicates. Statistics were assessed using an unpaired Student’s T-test (*, p < 0.05; **, p < 0.005). Error bars stand for the mean ± SD.

## MATERIALS AVAILABILITY

Plasmids generated in this study are available upon signing up MTA.

## DATA AND CODE AVAILABILITY

- The coordinates and EM density maps have been deposited to PDB and EMDB under accession codes of (9WGX, EMD-65961; 9WGY, EMD-65962; 20YQ, EMDB-67399) and are publicly available as of the date of publication.
- The protein mass spectrometry raw data have been deposited to the ProteomeXchange Consortium under accession code of PXD059713.
- Original images of uncropped gels are provided with this paper.
- Source data are provided with this paper.
- This paper does not report original code.
- Any additional information required to reanalyze the data reported in this paper is available from the corresponding authors upon request.

## AUTHOR CONTRIBUTIONS

LZM and XQ conceived the project. JX, YZ, XY, YD, JG, JD, WZ, and XL designed and performed the biochemical and cellular experiments and analyzed the data. HX collected Cryo-EM data and analyzed the data. JX and YZ curated data. SFS, ZL, XQ, and LZM supervised the experiments. XQ and LZM determined the structures and analyzed the data. LZM, XQ, JX, HX, YZ, and LZ drafted the manuscript.

## CONFLICTS OF INTEREST

LZM, XQ, JX, YZ, XY, JG, JD, and WZ are co-inventors on a patent application filed by Tianjin University relating to the work in this study.

## ACKNOWLEDGMENTS

The authors are grateful to Dr. Yufeng Yang at Fuzhou University for sharing the *PINK1*-KO COS7 cells. The authors are grateful to staff members of the Cryo-EM center (Southern University of Science and Technology) for their technical support. The authors are grateful to Drs. Yan Gao and Xiangyang Zhang at Tianjin University School of Pharmaceutical Science and Technology for their technical support on mass spectrometry. The authors are grateful to the financial support from National Natural Science Foundation of China to LZM (31670738, 31470730), XQ (31400645), and ZL (81870246, 82070329).

