## Supplemental Information for "MOLECULAR BASIS FOR PINK1 MATURATION"

### SUPPLEMENTARY FIGURES

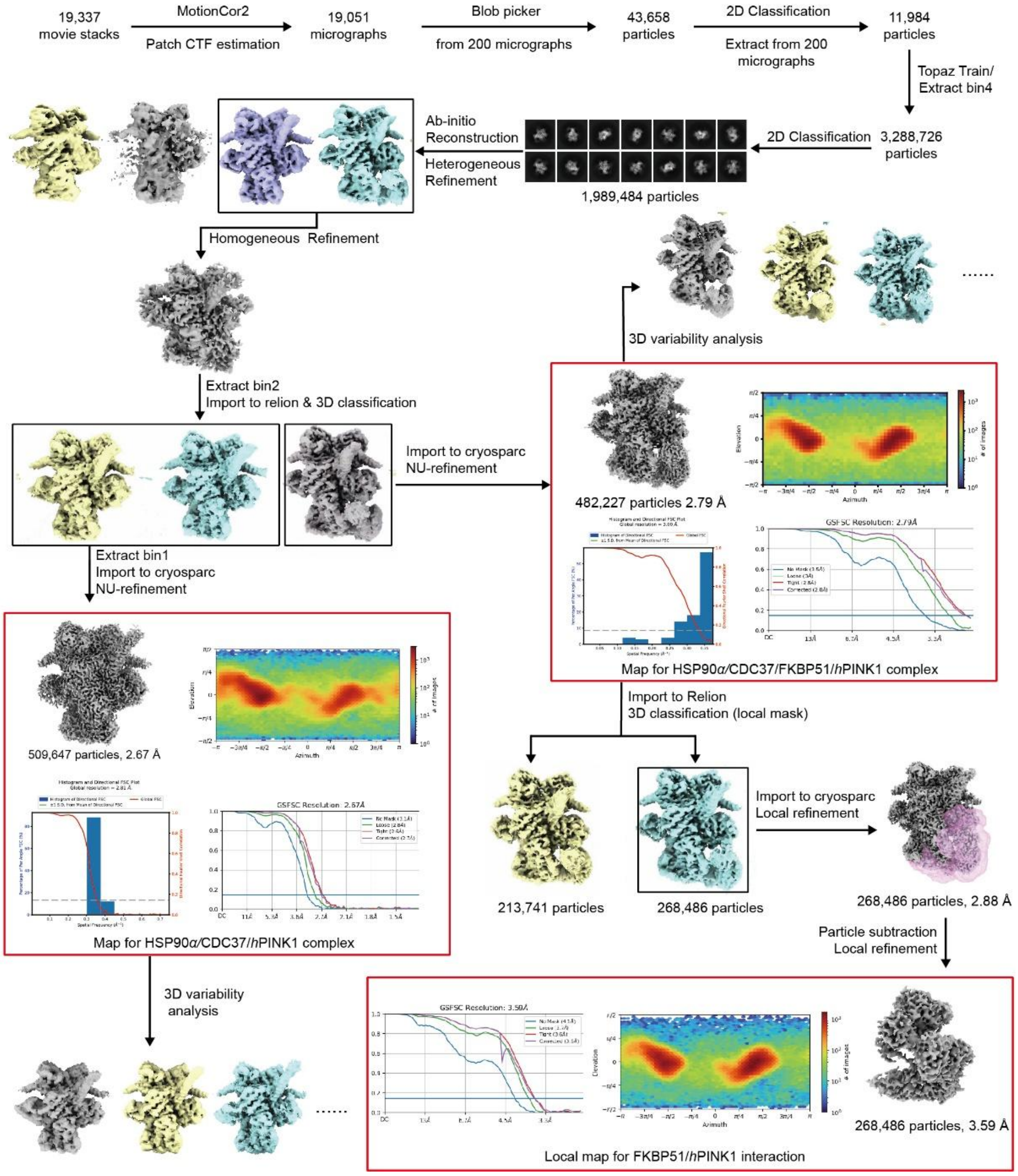


**Supplementary Fig. 1. Overall strategies and statistics in processing the Cryo-EM data of HSP90α/CDC37/PINK1 complexed with or without FKBP51.** Maps used for structural analysis are highlighted with red boxes.

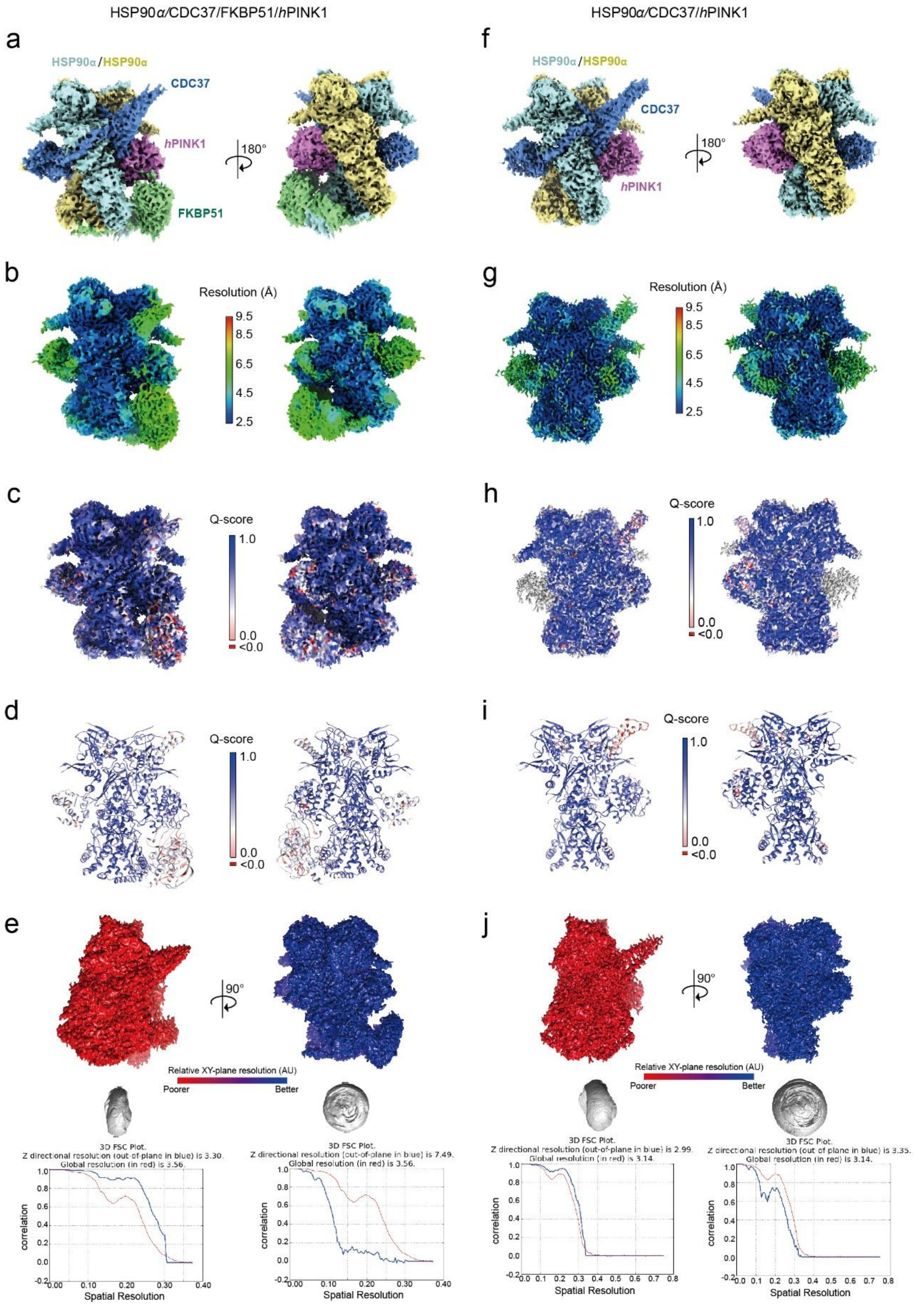


**Supplementary Fig. 2. Validation maps and models of the PINK1-chaperone complex structures. (a)** Cryo-EM density map of the HSP90α/CDC37/FKBP51/PINK1 complex contoured at 3σ, segmented and colored by protein component. **(b)** Density map of the HSP90α/CDC37/FKBP51/PINK1

complex colored by local resolution. **(c)** Density map of the HSP90α/CDC37/FKBP51/PINK1 complex colored by model Q-score. **(d)** Atomic model of the HSP90α/CDC37/FKBP51/PINK1 complex in cartoon representation, colored by model Q-score. **(e)** 3D FSC analysis of the HSP90α/CDC37/FKBP51/PINK1 complex. Top, density map colored by XY-plane resolution. Middle, 3D FSC volume displayed at the 0.143 threshold as an isosurface. Bottom, 3D FSC plot showing Z-directional (blue) and global (red) resolutions. **(f)** Cryo-EM density map of the HSP90α/CDC37/PINK1 complex contoured at 3σ, segmented and colored by protein component. **(g)** Local resolution map of the HSP90α/CDC37/PINK1 complex. **(h)** Density map of the HSP90α/CDC37/PINK1 complex colored by model Q-score. **(i)** Atomic model of the HSP90α/CDC37/PINK1 complex in cartoon representation, colored by Q-score. **(j)** 3D FSC analysis of the HSP90α/CDC37/PINK1 complex. Top, density map colored by XY-plane resolution. Middle, 3D FSC volume displayed at the 0.143 threshold as an isosurface. Bottom, 3D FSC plot showing Z-directional (blue) and global (red) resolutions.

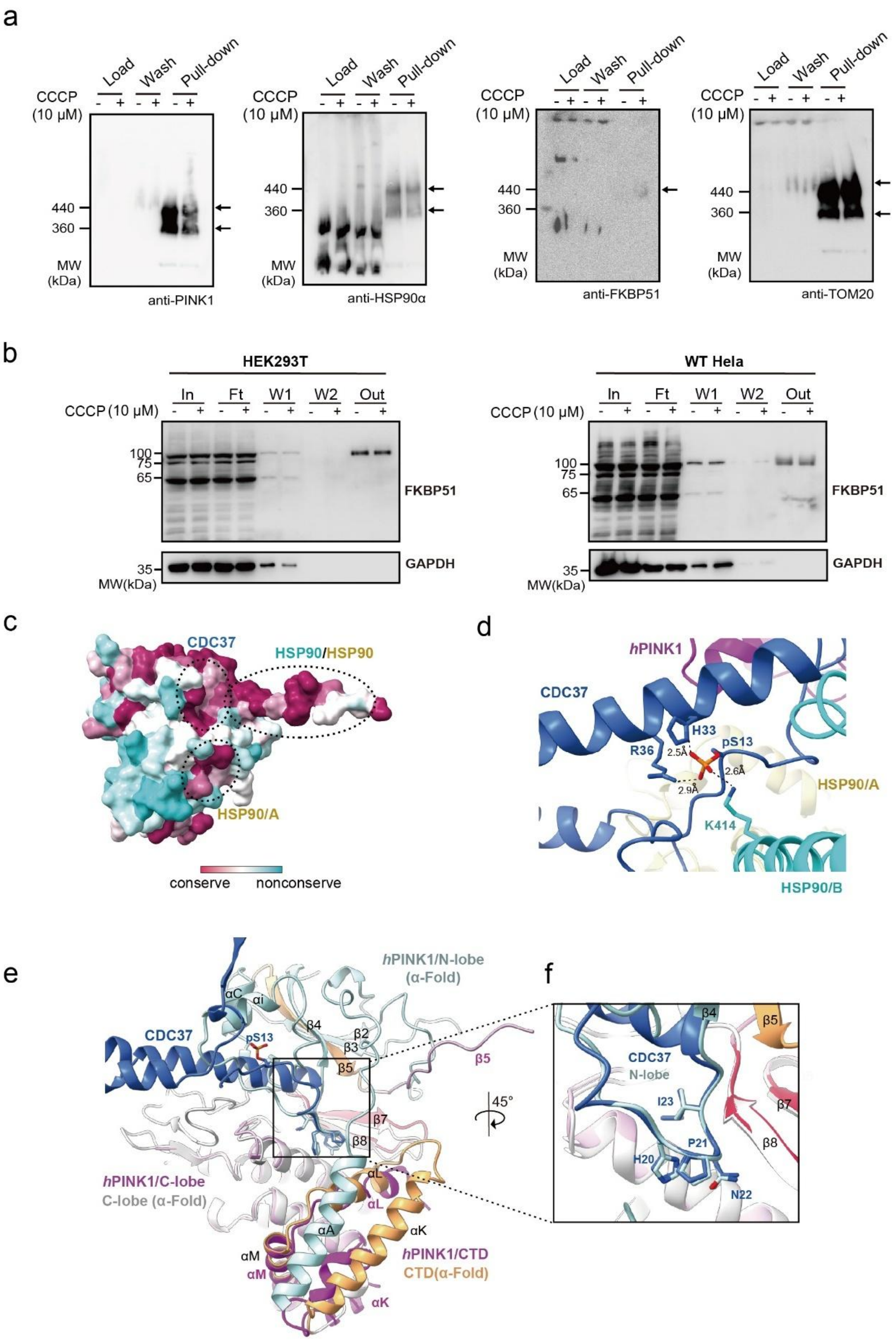


**Supplementary Fig. 3. Interaction of PINK1 with the HSP90α/CDC37 complex.** **(a)** Blue Native PAGE analysis of immunoprecipitated endogenous PINK1 complexes. WT Hela cells were treated with 10 μM CCCP or left untreated for 8 h.

PINK1-associated proteins were immunoprecipitated with an anti-PINK1 antibody and subjected to Blue Native PAGE. Bound proteins were analyzed by Western blotting using antibodies against PINK1, HSP90α, FKBP51, and TOM20. **(b)** Co-immunoprecipitation of endogenous PINK1-associated FKBP51. HEK293T and WT HeLa cells were treated with 10 μM CCCP or left untreated for 8 h. PINK1-associated FKBP51 was immunoprecipitated with anti-PINK1 antibody and analyzed by Western blotting using anti-FKBP51 antibody. SUMOylated FKBP51 was selectively enriched in the co-immunoprecipitation. In, input; FT, flowthrough; W1, wash 1; W2, wash 2; Out, pull-down. **(c)** Mapping of HSP90α- and CDC37-recognition sites on the PINK1 surface, colored by residue conservation (purple, conserved; cyan, non-conserved). **(d)** pSer13 of CDC37 interacts with CDC37 Arg36 and His33, as well as HSP90α Lys414, to stabilize the HPNI orientation. **(e)** Comparison of the AlphaFold2-predicted PINK1 structure with the chaperone-bound structure. The N-lobe of AlphaFold2-predicted PINK1 is shown in light blue cartoon representation, and the C-lobe in grey. The CTD and β5 strand are shown in gold. The chaperone-bound PINK1 is shown in pink, with the β7-β8 strands highlighted in red. CDC37 is shown in blue. **(f)** The HPNI motif of CDC37 mimics the αi-β4 loop of the PINK1 N-lobe, stabilizing the PINK1 C-lobe.

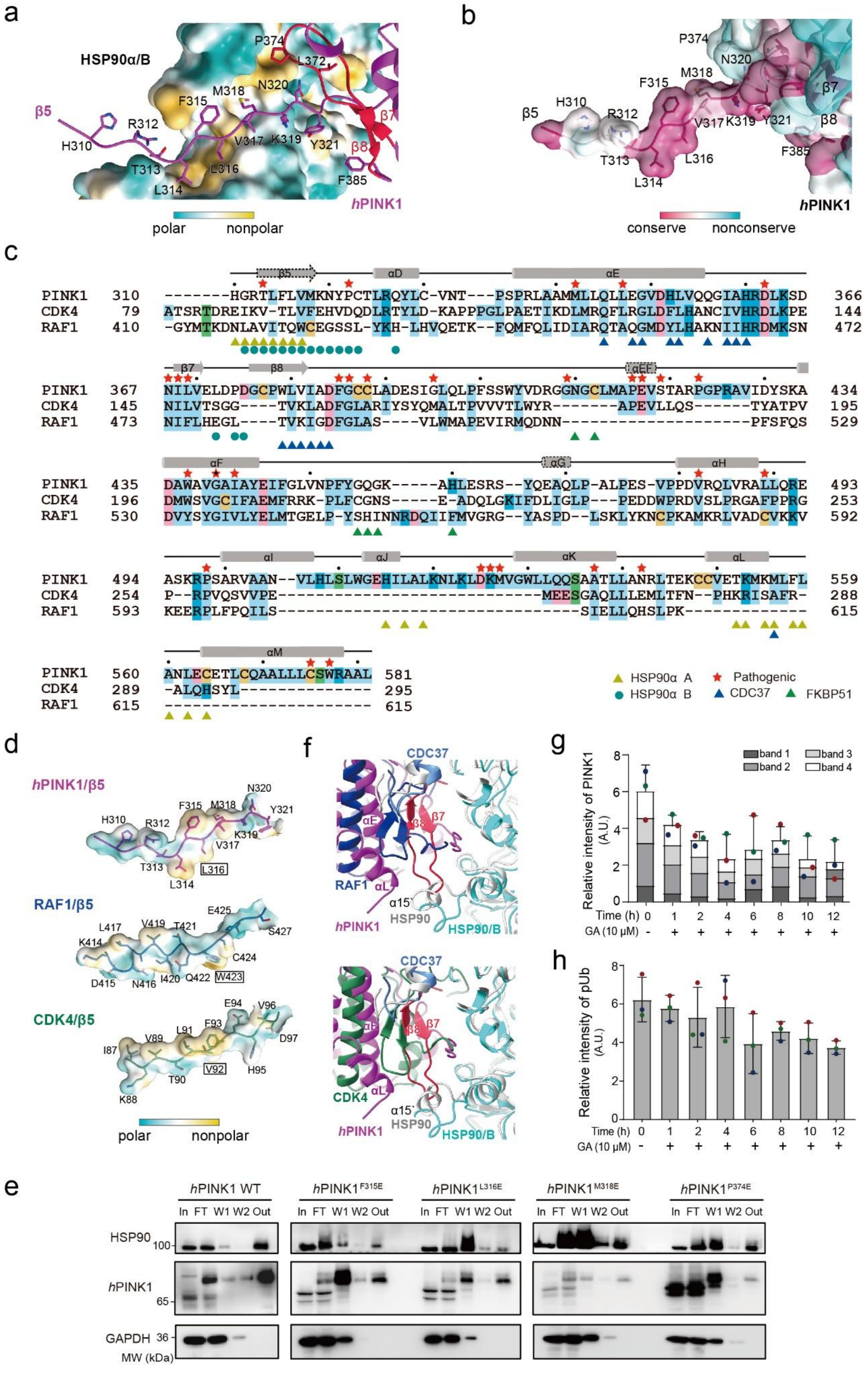


**Supplementary Fig. 4. Recognition of PINK1 by the HSP90α/CDC37/FKBP51**

**complex**. **(a)** The extended β5 strand of PINK1 fits into a hydrophobic tunnel of HSP90α. HSP90α subunit B is shown in surface representation, colored by hydrophobicity (cyan, polar; yellow, nonpolar). **(b)** Conservation of the PINK1 β5 strand. The PINK1 surface is colored by residue conservation. **(c)** Sequence alignment of HSP90/CDC37 kinase clients. Conserved residues are highlighted; every $10^{th}$ residue is marked with a black dot. α helices and β strands of PINK1 are shown as rods and arrows, respectively. Interface residues are indicated by symbols at the bottom; EOPD-associated mutations are marked with red stars. **(d)** Extended β5 strands from HSP90/CDC37 kinase clients shown in stick representation against surfaces colored by hydrophobicity. Central hydrophobic residues L316 (PINK1), W423 (RAF1), and V92 (CDK1) are aligned within the HSP90α lumen. **(e)** Mutations in the PINK1 β5 strand impair association with HSP90α. Plasmids encoding $PINK1^{WT}$, $PINK1^{F315E}$, $PINK1^{L316E}$, $PINK1^{M318E}$, and $PINK1^{P374E}$ were transiently transfected into *PINK1*-KO HeLa cells. HSP90α-bound proteins were immunoprecipitated from transfected cells using a specific antibody. Bound proteins were separated by SDS-PAGE and analyzed by Western blotting with antibodies against HSP90α, PINK1, and GAPDH. In, input; FT, flowthrough; W1, wash 1; W2, wash 2; Out, pull-down. **(f)** Structural comparison of PINK1 (magenta) with RAF1 (blue, top) and CDK4 (green, bottom), superimposed on their respective HSP90 complexes. The disordered α15` of HSP90α moves away to accommodate the projecting β7-β8 loop of PINK1. The HSP90/CDC37 complex with RAF1 or CDK4 is shown in grey, whereas the complex with PINK1 is colored as follows: HSP90α subunit B in cyan, CDC37 in slate. **(g,h)** Statistical analysis of Western blotting results of geldanamycin effects on PINK1 degradation (g) and Ub phosphorylation activity (h).

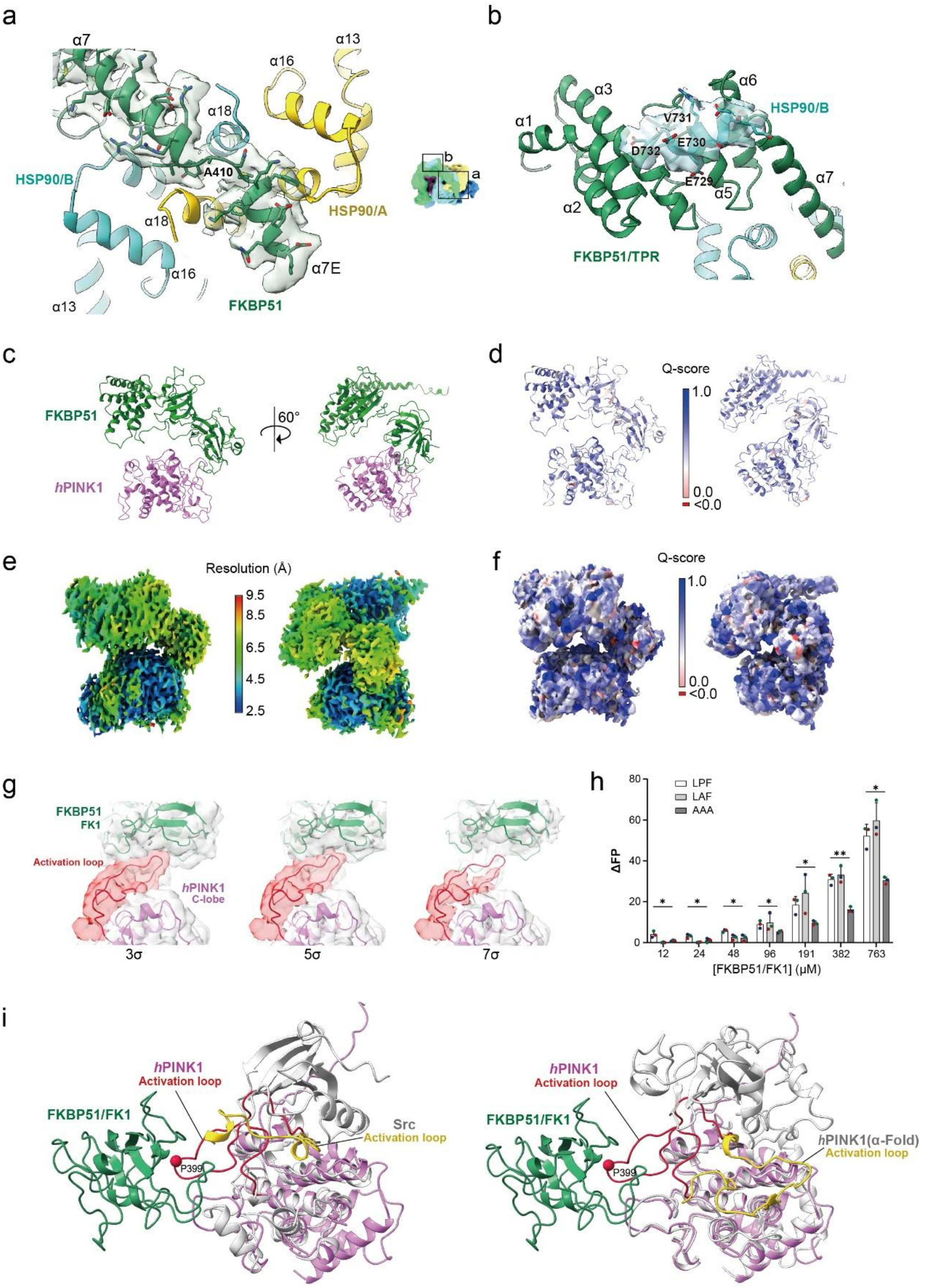


**Supplementary Fig. 5. Interaction of HSP90α-bound FKBP51 with inactive PINK1 activation loop. (a)** The α7E helix of FKBP51 interacts with the hydrophobic groove of HSP90α at the C-terminal dimeric interface. Density for the α7E helix is contoured at 3σ and shown as a green transparent surface; the helix is broken at A410. **(b)** The MEEVD motif of HSP90α binds within a cavity of the FKBP51 TPR domain, with density shown as a cyan transparent surface. **(c-g)** Validation maps and models for the locally refined FKBP51/PINK1 structure. **c** Cartoon representation of the FKBP51/PINK1 complex. FKBP51 is colored green and PINK1 pink. **d** Cartoon representation of the FKBP51/PINK1 complex colored by model Q-score. **e** Locally refined Cryo-EM density map of the FKBP51/PINK1 complex contoured at 3σ, colored by local resolution. **f** Local density map of the FKBP51/PINK1 complex contoured at 3σ, colored by model Q-score. **g** Local density map of the FKBP51/PINK1 complex low-pass filtered to 5 Å and contoured at 3σ, 5σ, and 7σ. Density is shown as transparent grey; the model is shown in cartoon representation. FKBP51 is in green; PINK1 is in pink with the activation loop in red. **(h)** Binding analysis of the FK1 domain to peptides derived from PINK1 activation loop. Fluorescently labeled peptides derived from PINK1 activation loop and its mutants were titrated into FK1 domain solutions at the indicated concentrations. Fluorescent anisotropy was recorded and analyzed. Each point represents the mean of triplicate technical repeats; binding statistics were derived from triplicate biological repeats. **(i)** The FK1 domain of FKBP51 recognizes the inactive conformation of the PINK1 activation loop. Inactive Src kinase (PDB 7NG7) and AlphaFold2-predicted active PINK1 were superimposed on the PINK1-chaperone complex. Superimposed structures are shown in grey, with activation loops in gold. PINK1 in the chaperone complex is shown in pink, with the activation loop in red. P399 is shown as a red sphere.

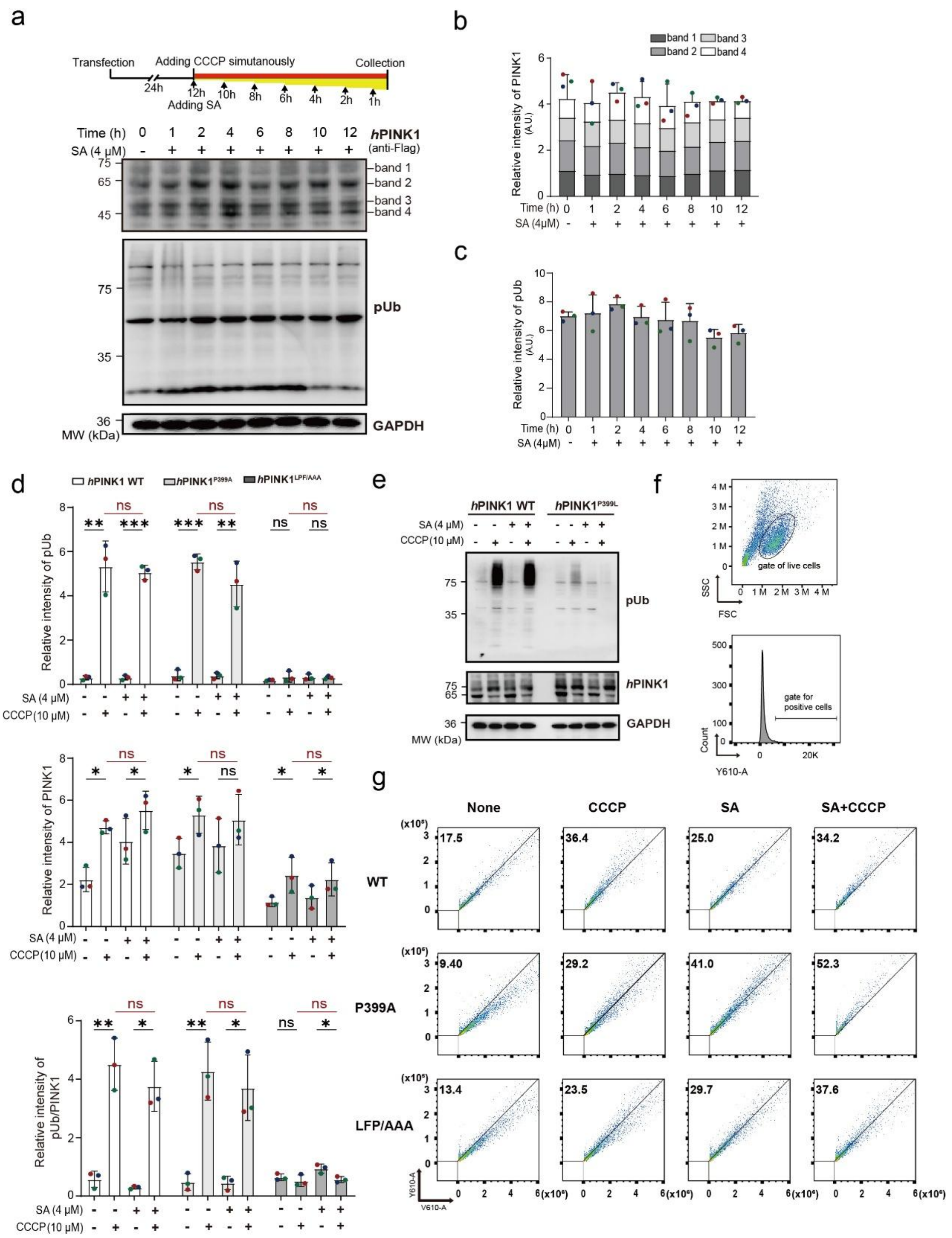


**Supplementary Fig. 6. Effects of SAFit2 on PINK1 stability and activity. (a-c)** Effects of SAFit2 on PINK1 stability and ubiquitin phosphorylation activity in transfected HEK293T cells. Cells were treated with 5 μM CCCP and 4 μM SAFit2 at the indicated time points. a, Representative Western blot. b,c, Statistical analysis from triplicate experiments. **(d)** Ubiquitin phosphorylation by PINK1 and mutants in transfected *PINK1*-KO COS7 cells treated with SAFit2 and/or CCCP. Ubiquitin phosphorylation, PINK1 expression and specific kinase activity were analyzed by

Western blotting. The statistics of triplicate biological repeats were determined by paired Student's T-test. **(e)** The P399L mutation impairs PINK1 kinase activity in Ubiquitin phosphorylation, as detected by Western blotting. **(f)** Gating strategy for flow cytometry analysis of PINK1-mediated mitophagy. Gates for live cells (top) and mt-Keima-positive cells (bottom). **(g)** Mitophagy in cells expressing $PINK1^{WT}$, $PINK1^{P399A}$ and $PINK1^{LPF/AAA}$ treated with SAFit2 and/or CCCP, analyzed by flow cytometry. Representative results from triplicate biological repeats.

#### Supplementary Table 1. Cryo-EM data collection, refinement and validation statistics

| | HSP90α/CDC37/PINK1/FKBP51 (EMDB:65961) (PDB: 9WGX) | HSP90/CDC37/PINK1 (EMDB:65962) (PDB: 9WGY) | PINK1/FKBP51 (EMDB:67399) (PDB: 20YQ) |
|---|---|---|---|
| **Data collection and processing** | | | |
| Microscope | FEI Titan Krios | FEI Titan Krios | FEI Titan Krios |
| Voltage (kV) | 300 | 300 | 300 |
| Detector | Gatan K3 Summit | Gatan K3 Summit | Gatan K3 Summit |
| Magnification | 130,000 × | 130,000 × | 130,000 × |
| Pixel size (Å) | 0.668 | 0.668 | 0.668 |
| Electron exposure ($e^{-}$/Å$^2$) | 60 | 60 | 60 |
| Defocus range (μm) | -1.2 to -2.0 | -1.2 to -2.0 | -1.2 to -2.0 |
| Automation software | EPU | EPU | EPU |
| Energy filter slit width (eV) | 20 | 20 | 20 |
| Micrographs used (no.) | 19,337 | 19,337 | 19,337 |
| Initial particle images (no.) | 3,288,726 | 3,288,726 | 3,288,726 |
| Final particle images (no.) | 482,227 | 509,647 | 509,647 |
| Map resolution (Å) | 2.79 | 2.67 | 2.67 |
| FSC threshold | 0.143 | 0.143 | 0.143 |
| **Refinement** | | | |
| Initial model used | 5FWM | 5FWM | 5FWM |
| Model resolution (Å) | 3.04 | 2.6 | 3.59 |
| FSC threshold | 0.143 | 0.143 | 0.143 |
| Map sharpening *B* factor (Å$^2$) | -88.5 | -105.1 | -122.1 |
| **Model composition** | | | |
| Non-hydrogen atoms | 17521 | 13087 | 6567 |
| Protein residues | 2161 | 1607 | 838 |
| Ligands | 4 | 4 | 0 |
| *B* factors (Å$^2$) | | | |
| Protein | 110.79 | 42.36 | 39.99 |
| Ligand | 53.29 | 21.58 | -- |
| **R.m.s.deviations** | | | |
| Bond lengths (Å) | 0.002 | 0.002 | 0.002 |
| Bond angles (°) | 0.498 | 0.434 | 0.510 |
| **Validation** | | | |
| Molprobity score | 1.74 | 1.41 | 1.97 |
| Clash score | 7.42 | 3.78 | 10.04 |
| Rotamers outliers (%) | 0.00 | 0.00 | 0.00 |
| **Ramachandran plot** | | | |
| Favored (%) | 95.26 | 96.43 | 92.99 |
| Allowed (%) | 4.74 | 3.66 | 7.01 |
| Outliers (%) | 0.00 | 0.00 | 0.00 |

**Supplementary Video 1. Conservation analysis of PINK1 interactions with HSP90α/CDC37/FKBP51.** This animation presents the 3D structure of the *h*PINK1 C-lobe in surface representation, colored by conservation levels (from cyan for non-conserved to maroon for conserved residues). FKBP51 is depicted in green, while HSP90α/CDC37 is shown in light yellow.

**Supplementary Video 2. Dynamic** c**onformations of the HSP90α/CDC37/*h*PINK1** c**omplex.** This animation displays classified electron density maps for 20 conformational states of the HSP90α/CDC37/hPINK1 complex. The density maps are contoured at 3 σ. The dynamic changes of the complex are shown across 20 consecutive frames, with subunits colored as in Fig. 1A.

**Supplementary Video 3. Morphed dynamic conformational transition of *h*PINK1.** This video illustrates the transition of *h*PINK1 between its unfolded state within the HSP90α/CDC37/FKBP51/*h*PINK1 complex and its folded state predicted by AlphaFold2. The morphing was performed in Chimera. Structural models are aligned at the kinase C-lobes and displayed in cartoons. The N-lobe and C-lobe of the kinase are colored in gray and magenta, respectively. Significant conformational changes are highlighted, with the β5 strand and C-terminal domain (CTD) shown in gold. The subunits of HSP90α, CDC37, and FKBP51 in the complex are colored as in Fig. 1A.

**Supplementary Video 4. Dynamic** c**onformational** c**hanges of the HSP90α/CDC37/FKBP51/*h*PINK1** c**omplex.** This movie illustrates classified electron density maps for the 20 conformational states of the HSP90α/CDC37/FKBP51/hPINK1 complex. The density maps are contoured at 3 σ. The dynamic changes are presented across 20 consecutive frames, with subunits colored as in Fig. 1A.
